# DeepDynaForecast: Phylogenetic-informed graph deep learning for epidemic transmission dynamic prediction

**DOI:** 10.1101/2023.07.17.549268

**Authors:** Chaoyue Sun, Ruogu Fang, Marco Salemi, Mattia Prosperi, Brittany Rife Magalis

## Abstract

In the midst of an outbreak or sustained epidemic, reliable prediction of transmission risks and patterns of spread is critical to inform public health programs. Projections of growth or decline among specific risk groups can aid in optimizing interventions, particularly when resources are limited. Phylogenetic trees have been widely used in the detection of transmission chains and high-risk populations. Moreover, tree topology and the incorporation of population parameters (phylodynamics) can be useful to reconstruct the evolutionary dynamics of an epidemic across space and time among individuals. We now demonstrate the utility of phylodynamic trees for infection forecasting in addition to backtracking, developing a phylogeny-based deep learning system, called *DeepDynaForecast*. Our approach leverages a primal-dual graph learning structure with shortcut multi-layer aggregation, and it is suited for the early identification and prediction of transmission dynamics in emerging high-risk groups. We demonstrate the accuracy of *DeepDynaForecast* using simulated outbreak data and the utility of the learned model using empirical, large-scale data from the human immunodeficiency virus epidemic in Florida between 2012 and 2020. Our framework is available as open-source software (MIT license) at: https://github.com/lab-smile/DeepDynaForcast.

**Author Summary:** During an outbreak or sustained epidemic, accurate prediction of patterns in transmission risk can reliably inform public health strategies. Projections indicating growth or decline of transmission for specific risk groups can significantly enhance the optimization of interventions, especially when resources are limited. To address this, we present *DeepDynaForecast*, a cutting-edge deep learning algorithm designed for forecasting pathogen transmission dynamics. Uniquely, *DeepDynaForecast* was trained on in-depth simulation data and used more information from the phylogenetic tree of pathogen sequence data than any other algorithm in the field to date, allowing classification of samples according to their dynamics (growth, static, or decline) with incredible accuracy. We evaluated the model’s performance using both simulated outbreak data and empirical, large-scale data from the HIV epidemic in Florida between 2012 and 2020. We conclude *DeepDynaForecast* represents a significant advancement in genomics-mediated pathogen transmission characterization and has the potential to catalyze new research directions within virology, molecular biology, and public health.

## Introduction

Epidemiological modeling of the spread of a disease during an outbreak is critical in predicting the devastation caused by the root pathogen and in designing specific public health interventions to curb unfavorable outcomes. The vast majority of the models used, however, often assume random mixing of the host population, which can affect parameter estimates used in these predictions (e.g., (1–3)). Awareness of this limitation often exists, but incorporation of the numerous relevant structures within a population requires specific *a priori* information regarding not only population behavior but also transmission routes of the pathogen responsible for the outbreak. For example, increased transmission of food-borne pathogens may be found among individuals primarily relying on a local food vendor (4). Moreover, the population structure may be dynamic, as in the case of brief social gatherings wherein airborne pathogens are more easily spread. Groups of infected individuals for which risk of pathogen transmission is heightened, regardless of their episodic nature, can be identified through combined pathogen molecular data and patient information (e.g., contact tracing). Identifying these groups can provide invaluable insight into specific patterns of spread for which targeted interventions can aid in curbing an outbreak.

Pathogen genomic data became increasingly used in outbreak surveillance primarily owing to the evolutionary information they provide for the development of therapeutic interventions, a primary recent example being SARS-CoV-2 (5). The evolutionary trajectory of a pathogen can be readily monitored owing to the rapid accumulation of mutations that result from the short generation time and/or infidelity of the replication machinery characteristic of micro-organisms such as viruses and bacteria. The rate of accumulation of mutations, particularly for viruses (6), is often deterministic and so is proportional to the number of replication cycles. Assuming the replication rate is relatively constant for a particular pathogen during infection, the number of replication cycles, and thus mutations, can be estimated from the time of infection of one individual and time of transmission from that individual to another. Fewer mutations are thus expected to occur for shorter transmission times when transmission is more likely. This relationship of evolution and transmission is the principle for genetic clustering, wherein in a population sample, transmission clusters are defined as groups of patient-derived pathogen sequences characterized by minimal genetic variation, representing high-risk transmission. Genetic clustering can be achieved using a variety of algorithms. Primarily distance-based methods (7, 8) rely on a user-specified threshold during pairwise genetic distance estimation among samples, below which samples are considered to be connected. These methods are fast and accurate; however, the resolution of these methods is limited to the level of the group, as specific mutation information is not used to resolve individual relationships. Alternatively, a Phylogenetic tree reconstructed from the set of mutational information at genomic sites offers increased resolution for individual relationships within the group. Moreover, branches within the tree can adopt generalized shapes, or topological features, that can provide critical information as to the underlying contact dynamic (9, 10) (e.g., presence of “super-spreaders”) and population dynamics over time (11). Integrating population genetic-parameterized methods with phylogenetics is known as ‘phylodynamics’ and is based on retroactively tracing an epidemic backward in time. Little is known, however, regarding the *predictive* power of phylodynamic information, particularly when supplied at the onset of an outbreak when samples are scarce.

In recent years, deep learning has been applied extensively in epidemiology, allowing for increased information gained from large datasets comprising numerous multi-factorial risk factors of infection and spread (12, 13). Neural networks have also gained popularity among sequencing endeavors, used in identifying connections between mutations within high-dimensional genome sequence data and disease status (14). Graph neural networks (GNN) have been specifically designed for data that could be represented as a graph structure connected via complex relationships (15–17), for which the phylogenetic tree is a prime candidate. Complex relationships within outbreak phylogenies can range from simple community-based structures, with isolated risk groups transmitting at constant rates, to risk group mixing with varying, non-deterministic transmission rates between groups and over time (Supplementary Figure 1). While previous research has attempted to predict infection dynamics using convolutional neural networks with sequencing data (18), these methods fall short in directly incorporating the hierarchical relationships among sequenced individuals. We explored this complexity by simulating phylogenies representing viral and bacterial outbreaks at varying times during the epidemic and comprised of smaller risk groups with deterministic, yet varying transmission rates relative to the background population. Using these data, we demonstrate the effectiveness of our GNN method (*DeepDynaForecast*) in predicting future dynamics from limited, early sampling and its promising capacity to improve public health efforts in preventing epidemic spread. We then sought to apply the trained model to the human immunodeficiency virus (HIV) epidemic in Florida between 2012 and 2020, for which large-scale pathogen sequencing data and metadata is available, previously analyzed using traditional distance-based methods (19).

## Results

### *DeepDynaForecast* framework

Our *DeepDynaForecast* model is developed to predict near-future transmission dynamics for the external nodes (leaves) on a Phylogenetic tree. Figure 1 illustrates the complete computational framework, composed of two primary components: a primal-dual graph learning architecture and a cross-layer dynamics prediction module. The detailed methodology of the framework is provided in the Methods section, while the ensuing sections provide succinct introductions to each key component.

**Fig 1.**
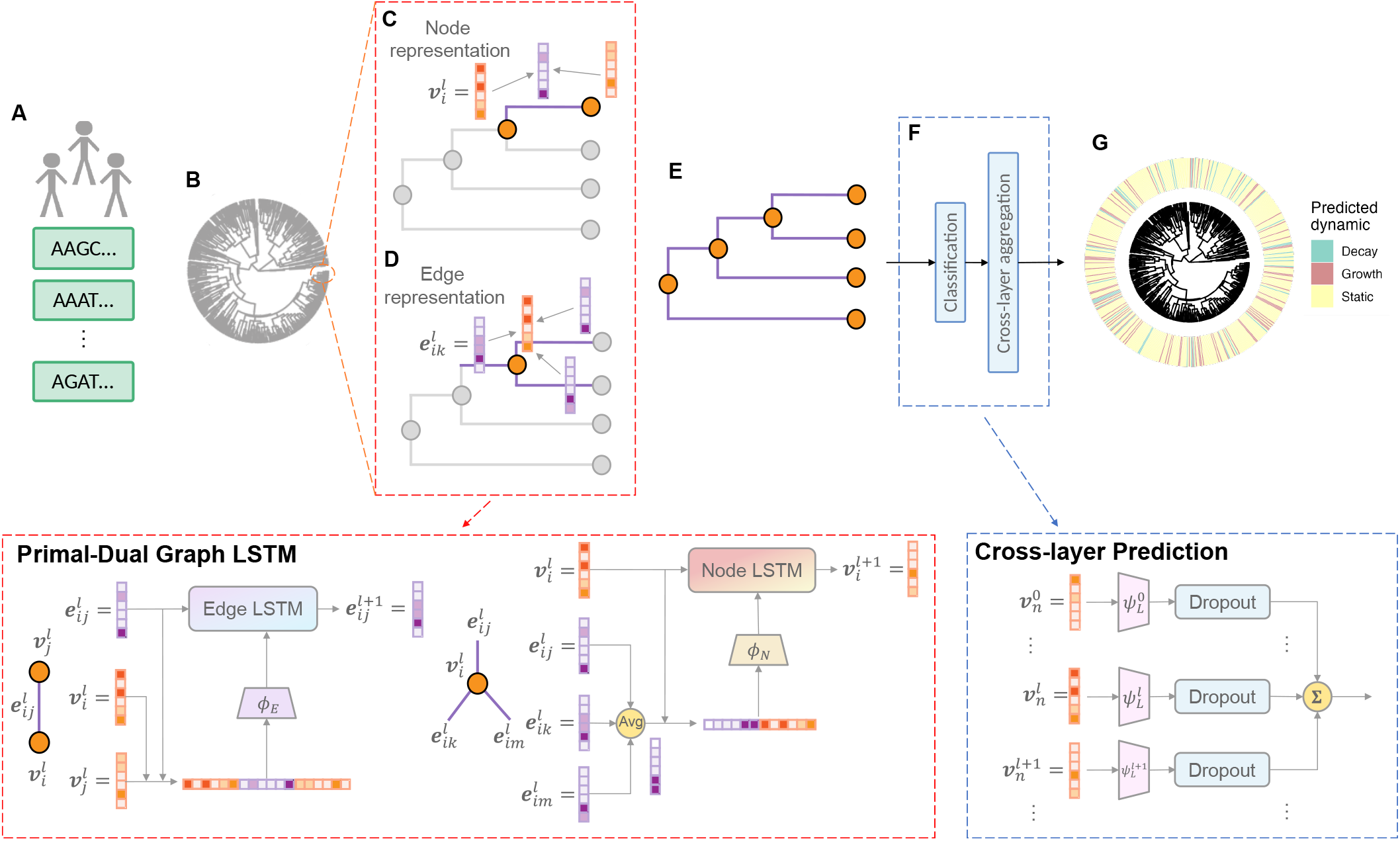
Our proposed *DeepDynaForecast* architecture. **A**. Pathogen genomic data collected during an outbreak. **B**. A Phylogenetic tree, reconstructed from the mutational information at genomic sites, was used to trace the transmission among the populations. In this tree, nodes represented individuals, while edges represented transmission or mutation events. We modeled the Phylogenetic tree using a bi-directed graph, where initial node representation vector **v**_*i*_ was randomly generated for each node *i*, and edge representation vector **e**_*ij*_ for edge *e*_*ij*_ from node *i* to *j* was initialized from the branch length with a neural network. **C-D**. Example of Primal-Dual Graph Long Short-Term Memory (PDGLSTM) learning architecture on a subtree to update **v**_*i*_ and **e**_*ij*_ at the *l*-th layer. Two parallel LSTM modules were utilized to update the node and edge representations in each message-passing iteration. Within this process, each edge/node aggregated adjacent node/edge representations and encoded low-dimensional messages by the neural networks *ϕ*_*E*_ and *ϕ*_*N*_. These node/edge messages were input into their corresponding LSTM modules to facilitate the update of node and edge representations **E**. This system sequentially applied *N* rounds of message-passing iterations, thus producing updated nodes and edges representations. **F**. Cross-layer Prediction (CLP) module on each leaf node. A series of neural networks *{ψ*_*L*_*}* were engaged in predicting the dynamics of leaf *n* using various levels of node representations *{***v**_*n*_*}*. This process was followed by dropout layers and summation operations to generate the final prediction. **G**. Predicted dynamics for leaves on the Phylogenetic tree.

### Problem formulation

The Phylogenetic tree, generated from the transmission networks within *nosoi* (20) agent-based stochastic simulation platform, is modeled as a bi-directed graph, with initial node representation vectors **v** randomly generated and edge representation vectors **e** initialized from the branch length via a neural network. The nodes within the tree are grouped into two categories: 1) background infected population, representing transmission among the majority of infected individuals, and 2) risk groups composed of differing transmission dynamics. The nodes belonging to risk groups are categorized into three distinct groups -static, growth, or decay -based on pre-defined transmission dynamics during the simulation (Supplementary Table 1). These categories were used as classification labels during our supervised learning modeling.

### Primal-dual graph LSTM module (PDGLSTM)

The PDGLSTM (21, 22) learns the node and edge representations simultaneously by applying two parallel GLSTMs on nodes and edges. GLSTM incorporates LSTM to learn about the sequential representations generated from multiple message-passing iterations. As shown in Figure 1, each edge/node aggregates neighboring node/edge representations and encodes low-dimensional messages by the neural networks *ϕ*_*E*_ and *ϕ*_*N*_. These node/edge messages are input into their corresponding LSTM modules to facilitate the update of node and edge representations.

### Cross-layer Prediction module (CLP)

After sequentially *N* times of message-passing iterations, the CLP module is applied to predict the dynamics of leaves and masks internal nodes when predicting. A series of neural networks {*ψ*_*L*_} are engaged in predicting the dynamics of leaves using various levels of node representations {**v**}. This process is followed by dropout layers and summation operations to make the final result robust.

### *DeepDynaForecast* performance on simulated outbreaks

This section comprehensively evaluates DeepDynaForecast’s performance in predicting the dynamic status. Table 1 compares two baseline models -Graph Convolutional Network (GCN) (23) and Graph Isomorphism Network (GIN) (15) - and DeepDynaForecast models in three scenarios: training on acute infectious respiratory virus (ARI) simulations only (DDF_ARI), *Mycobacterium Tuberculosis* (TB) simulations only (DDF_TB), and their combination (DDF_ARI_TB). Two baseline models were trained on the combined simulations and all five models were evaluated on ARI, TB, and their combination in Table 1. The metrics included accuracy, F1-score, precision, area under the receiver operating characteristic (AUROC), Brier score (BS), and Cross-entropy (For further details, see performance evaluation in the Methods section).

**Table 1.** Performance for two baseline models and *DeepDynaForecast* on Transmission Dynamics Prediction of External Nodes with Three Training Scenarios. Baseline models comprised Graph Convolutional Network (GCN) and Graph Isomorphism Network (GIN). The training scenarios included the model trained with Respiratory virus simulations only (DDF_ARI), the model trained on *Mycobacterium tuberculosis* simulations only (DDF_TB) and the model trained on the mixed type of transmission patterns (DDF_ARI_TB). Baseline models were also trained on the combination of ARI and TB. To mitigate the impact of unbalanced label distribution, metrics for multi-class node types—including accuracy, F1-score, precision, and area under the receiver operating characteristic (AUROC)—were uniformly aggregated across the three classes. Weighted Brier score (BS) and weighted Cross-entropy (CE) were calculated based on predicted probabilities adjusted by the inverse prevalence of classes, providing “soft” evaluations of the models. For each testing dataset, models were assessed on ARI trees, TB trees, and a combination of both. The best performance for each evaluation metric among the five models is highlighted using underlining, italics, and bold for ARI, TB, and the combination, respectively.

| Model | Dataset | Balanced Accuracy ↑ | Macro F1 ↑ | Macro Precision ↑ | Macro AUROC ↑ | Weighted BS ↓ | Weighted CE ↓ |
| --- | --- | --- | --- | --- | --- | --- | --- |
| GCN | ARI | 0.640 | 0.298 | 0.370 | 0.753 | 0.444 | 0.729 |
|  | TB | 0.556 | 0.240 | 0.350 | 0.676 | 0.537 | 0.899 |
|  | ARI+TB | 0.612 | 0.274 | 0.362 | 0.737 | 0.470 | 0.775 |
| GIN | ARI | 0.668 | 0.308 | 0.377 | 0.767 | 0.421 | 0.696 |
|  | TB | 0.585 | 0.284 | 0.354 | 0.710 | 0.528 | 0.894 |
|  | ARI+TB | 0.654 | 0.299 | 0.368 | 0.763 | 0.442 | 0.736 |
| DDF_ARI | ARI | 0.865 | 0.480 | 0.436 | 0.960 | 0.201 | 0.369 |
|  | TB | 0.694 | 0.268 | 0.365 | 0.853 | 0.443 | 0.791 |
|  | ARI+TB | 0.800 | 0.367 | 0.387 | 0.918 | 0.293 | 0.528 |
| DDF_TB | ARI | 0.627 | 0.487 | 0.597 | 0.886 | 0.636 | 1.622 |
|  | TB | 0.720 | 0.440 | 0.404 | 0.943 | 0.393 | 0.695 |
|  | ARI+TB | 0.627 | 0.472 | 0.432 | 0.862 | 0.626 | 1.557 |
| DDF_ARI_TB | ARI | 0.895 | 0.494 | 0.452 | 0.969 | 0.157 | 0.284 |
|  | TB | 0.744 | 0.392 | 0.381 | 0.925 | 0.345 | 0.604 |
|  | ARI+TB | 0.881 | 0.459 | 0.424 | 0.964 | 0.175 | 0.315 |

For the model performance on combined ARI and TB datasets, DDF_ARI_TB achieved 88.1% balanced accuracy and 0.964 macro AUROC, outperforming the best baseline model (GIN) by 34.7% and 26.3%. Another significant improvement from GIN to DDF_ARI_TB is observed in the “soft” evaluations of the model. For example, DDF_ARI_TB reduces the weighted Brier score by 60.4% (from 0.442 to 0.175) and shows a similar trend for weight Cross-entropy (from 0.775 to 0.315). Similar conclusions could be drawn considering all six metrics on either ARI testing or TB testing with GCN, GIN, and DDF_ARI_TB. This analysis demonstrates that the *DeepDynaForecast* architecture can learn more comprehensive transmission patterns compared to GCN and GIN.

To explore the impact of different training scenarios, we compared *DeepDynaForecast*’s performance when trained on ARI only, TB only, and their combination. When tested on combined ARI and TB datasets, DDF_ARI_TB improved balanced accuracy by 10.1% over DDF_ARI and 40.5% over DDF_TB. Considering all six metrics, DDF_ARI_TB consistently outperformed DDF_ARI, while substantial improvements were observed in four metrics for DDF_TB. These results suggest that a generalized training scenario enhances the model’s performance in real-world applications. Notably, even in evaluations focused solely on ARI, DDF_ARI_TB surpassed DDF_ARI in five of the six metrics, further supporting the generalization capabilities of a model trained on mixed transmission types. In terms of performance on TB, although TB trees were not part of DDF_ARI’s training, it still achieved a 69.4% balanced accuracy and a 0.853 AU-ROC, with only a 6.7% and 7.8% decrease in balanced accuracy and AUROC, respectively, compared to DDF_ARI_TB.

These findings indicate that DeepDynaForecast possesses exceptional generalization abilities for unseen transmission patterns and excels in learning mixed transmission patterns. While the TB-only model (DDF_TB) performed well when evaluated exclusively on TB data (attaining a balanced accuracy of 72.0% and a macro AUROC of 0.943), it fell short in predicting ARI nodes. This discrepancy suggests that the branch patterns characteristic of chronic infection epidemics, such as TB, are distinct from those associated with acute infections, like ARI. The observed performance deficit could be ascribed to the reduced representation of TB decay nodes (51,419) compared to ARI decay nodes (589,119) -for more details on node representation, refer to the Methods section. This observation further elucidates why the GCN, GIN, and DDF_ARI_TB models yield less effective performance on TB testing than ARI testing.

In addition to comparing method performances for specific labels, we also utilized the resulting confusion matrices to summarize and illustrate detailed prediction distributions across different cluster types. Figure 2A displays the confusion matrices of various approaches, with each element normalized by the number of true labels summarized over each row for easy comparison. Consistent with the conclusions drawn from the metrics, DDF_ARI_TB achieved 88.9%, 84.6%, and 90.7% accuracy on decay, static, and growth nodes, respectively, outperforming all five models.

**Fig 2.**
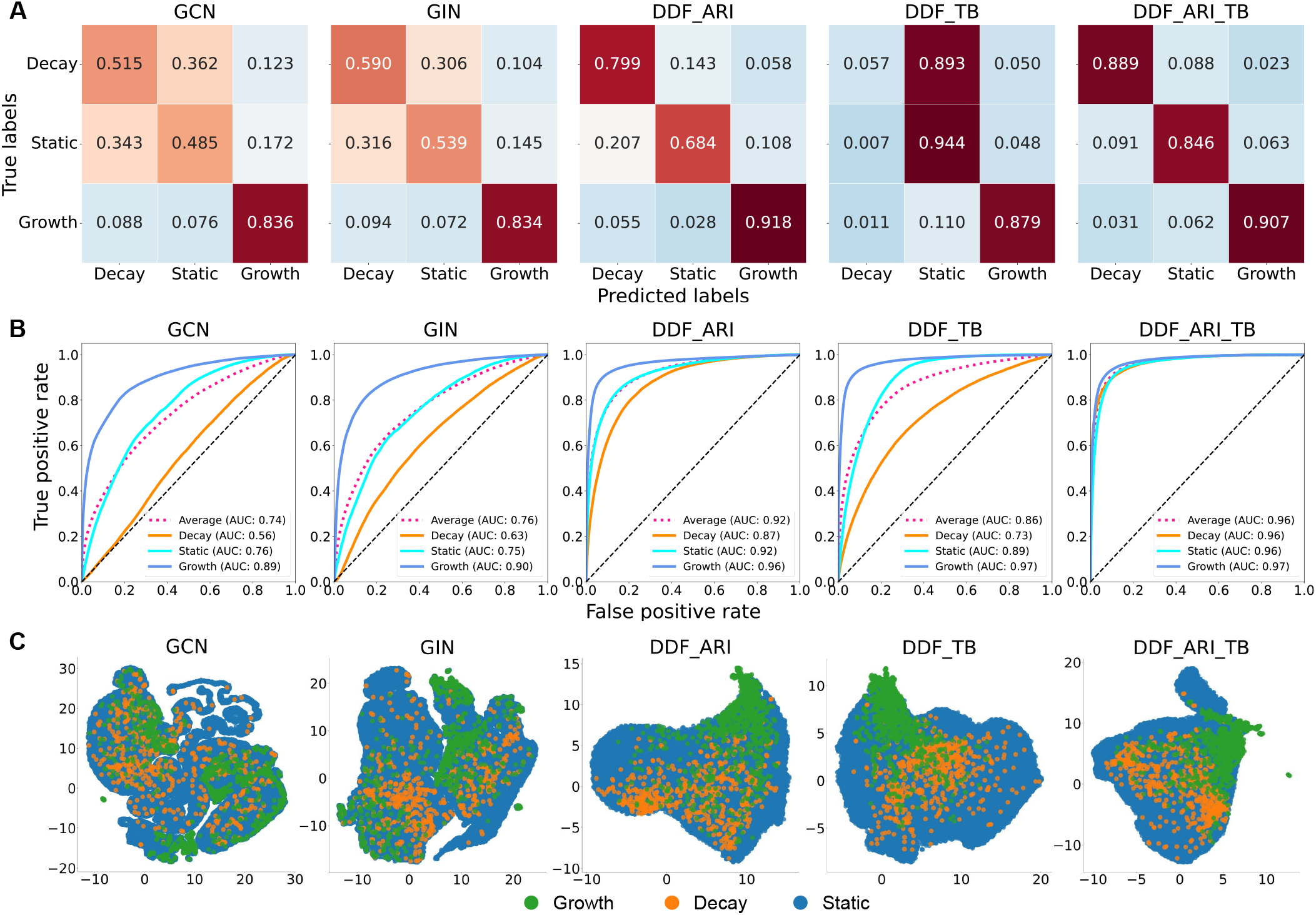
Figurative performance comparison of five models on combined ARI and TB test sets. **A**. Confusion matrices with row-wise normalized elements. **B**. One-verserest receiver operator characteristic curve (ROC) for each class and a macro averaged ROC curve with magenta dash lines. The corresponding AUCs are indicated for each curve. **C**. UMAP visualization of aggregation of learned node representations in message-passing iterations. Plots were generated in randomly sampled 50 Phylogenetic trees in ARI and TB test sets.

Decay and static leaves were often misclassified in GCN, GIN, and DDF_TB, but this issue was mitigated in DDF_ARI and DDF_TB. Figure 2B demonstrates one-versus-rest receiver operator characteristic curve (ROC) for each class and a macro-averaged ROC curve. DDF_ARI_TB attained *>* 0.96 AUROC for all four curves, exhibiting the best performance. Figure 2C presents the UMAP (24) 2-dimensional (2D) feature space generated from an aggregation of learned node representations. Decay nodes were less clustered in the UMAP space than growth samples, explaining why static and decay samples were more prone to misclassification. As expected, nodes with the same labels were better clustered in DDF approaches.

### Model Limitation Analysis

To assess the limitations of the *DeepDynaForecast* model for future application, we conducted limitation analysis to determine the predicting accuracy distribution over ground truth cluster size and risk groups. The results, presented in Figure 3, demonstrate a consistent reduction in accuracy across all leaves, decay leaves, and growth leaves for small cluster sizes. Our analysis also shows that the *DeepDynaForecast* model requires a sample size greater than 30 individuals to achieve the proposed level of performance. Furthermore, we evaluated the model’s performance on risk groups representing different transmission patterns. As shown in Figure 3B, the model achieved impressive performance for groups A (background leaves), F (growth leaves), and G (decay leaves) for ARI simulations. However, the model struggled to predict leaves in the rest groups representing static clusters accurately. This observation was consistent with the lower performance on cluster leaves in Supplementary Table 4, compared to background leaves and all leaves. Evaluations on TB draw similar conclusions, with impressive performance for groups A (background) and D (growth) but poor performance for groups B and C (static clusters). Additionally, poor performance in group E (decay) aligned with the findings of the confusion matrix. Overall, our analysis indicates limitations of the *DeepDynaForecast* model for future applications, particularly for smaller cluster sizes and certain risk groups.

**Fig 3.**
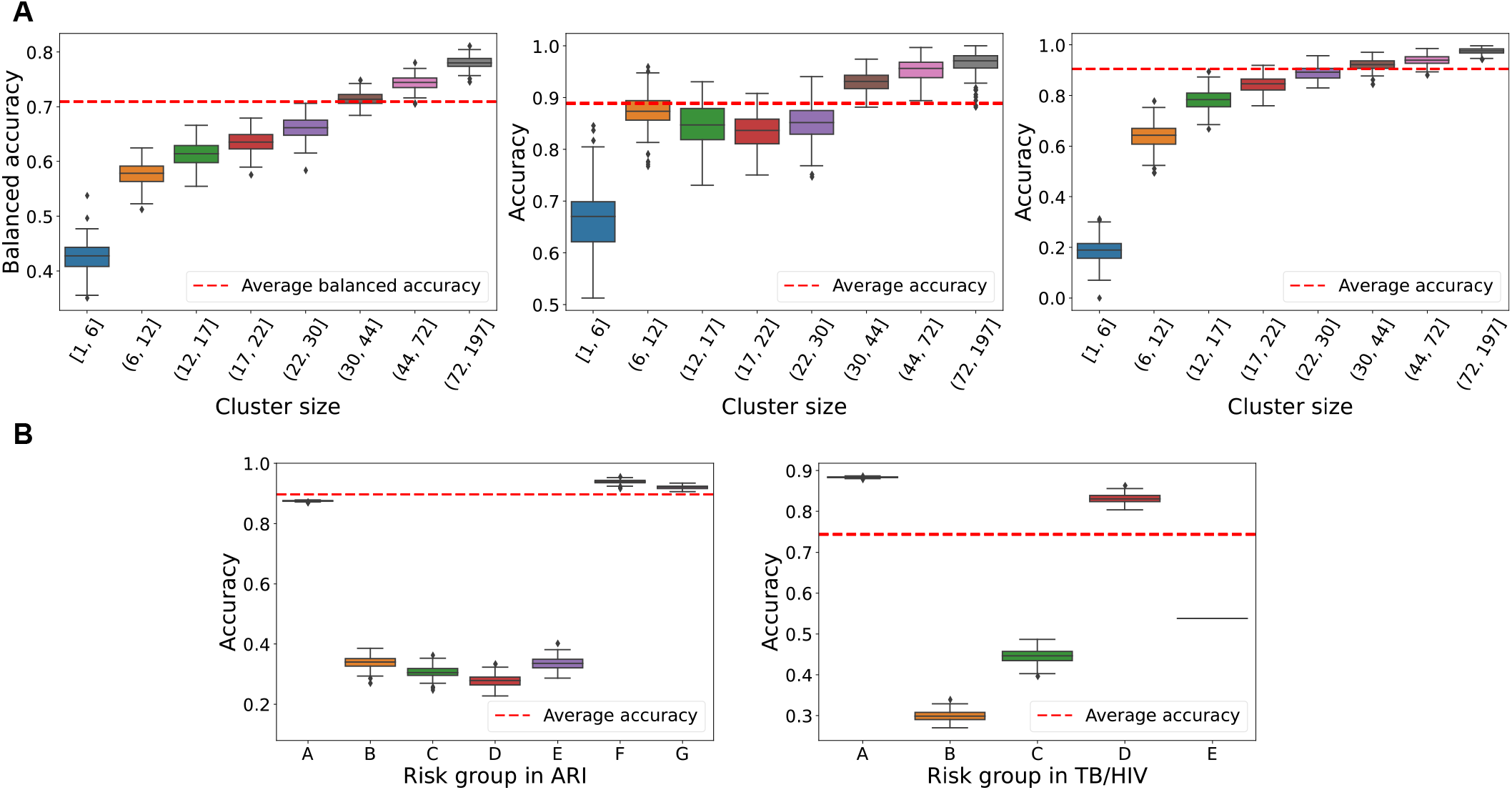
Sensitivity of the DeepDynaForecast model to cluster size and risk groups. **A**. Left panel: balanced accuracy on predicting all leaves with different external node sizes. Binned intervals for quantitative data were generated using eight quantiles. Model performance on decay clusters and growth clusters are illustrated in the middle and right panels, respectively. **B**. Performance of varying risk groups in ARI and TB simulations (see Supplementary Table 1).

### Transmission dynamics of the HIV epidemic in Florida

Persons living with HIV (PLWH) are an ideal example of a structured population with disproportionate transmission patterns across geographical regions and risk groups (25). In 2017, rates of new HIV diagnoses in the United States (U.S.) were highest in the southern region, where the state of Florida in particular exhibited the highest number of new diagnoses (25). In 2019, the U.S. Department of Health and Human Services released the federal plan for Ending the HIV Epidemic (EHE) within 10 years, identifying 48 counties with high incidences of HIV diagnoses, including seven urban Florida counties (Broward, Duval, Hillsborough, Miami-Dade, Orange, Palm Beach, and Pinellas) as high-priority areas. Strategic interventions for these areas largely revolved around building the capacity to detect and respond to ongoing and emerging clusters (26), dependent on molecular epidemiology techniques. The Florida Department of Health (FDOH) has been actively collecting partial HIV-1 polymerase (*pol*) genetic sequences from surveillance laboratories since 2007, to attain an analyzable HIV nucleotide sequence within 12 months of diagnosis for *>* 60% of persons diagnosed with HIV per year, culminating in a total of 28, 098 partial HIV-1 *pol* sequences. Of these, 27, 115 (96.5%) were classified as subtype B and used in previous clustering analysis by Rich *et al*. (19). In this study, MicrobeTrace (27) was used to identify potential clusters of transmission, wherein a genetic distance of 1.5% was used as a threshold, below which individuals were considered to be related via transmission chain or high-risk transmission group (Figure 4). Transmission clusters were then assessed for risk factors using patient meta-data provided from the FDOH, which included year of HIV genotype determination, year of birth, birth region, gender at birth, race/ethnicity, county of residence, and mode of transmission exposure. Mode of transmission exposure categories included men who have sex with men (MSM), heterosexual (HET), intravenous drug use (IDU), mother-to-child (MTC), and unknown transmission. The majority of the sequences in this study originated from metropolitan regions containing the highest HIV prevalence, including all seven recently identified EHE priority counties. It is important to note that the proportion of clustered sequences by county was not dependent on the number of sequences available (i.e., increased sampling did not result in a proportional increase in clustering). However, rural and suburban counties not considered EHE priority, with relatively high HIV prevalence, demonstrated a high proportion of unclustered sequences, indicating a relative abundance of missing infection links during surveillance. Sociodemographic and geographic characteristics were associated with a different propensity to cluster. In particular, age, gender, race/ethnicity, rural vs. urban residence, country of origin, mode of HIV transmission, all distributed differently among clustered and unclustered individuals, and across cluster sizes. The large clusters were also minimally assortative by age and sampling year, compared to smaller clusters (2-10 individuals). Regardless of cluster size, assortativity was highest for geographic region (i.e., county). It was duly noted that these associations should not be regarded as causal, because they could be confounded by other factors or due to selection bias, leading to wrong interpretations.

**Fig 4.**
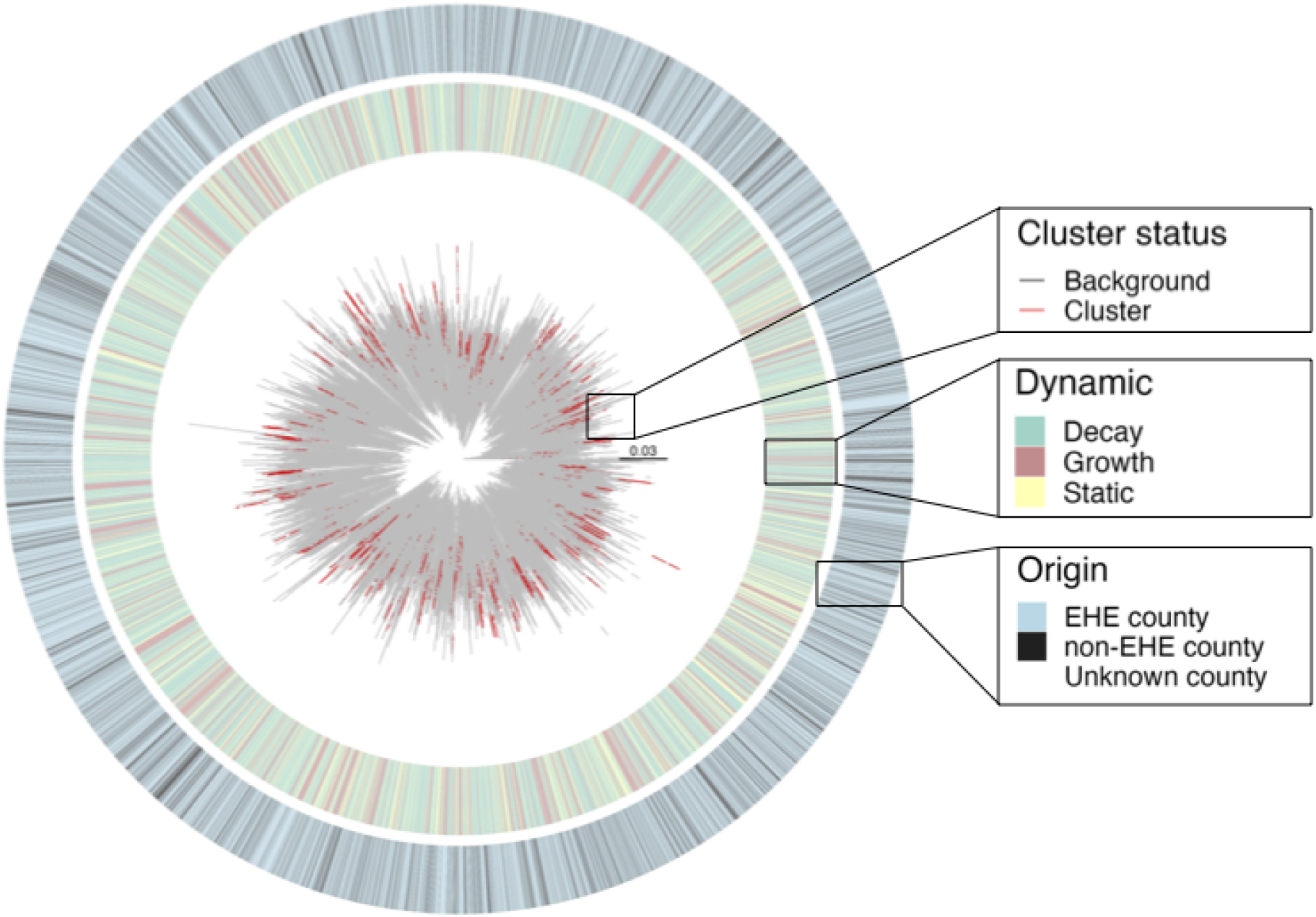
Florida HIV-1 subtype B *pol* sequence phylogeny (2012-2017). The maximum likelihood phylogeny was generated as described in Rich *et al*. (19) for 27, 115 partial *pol* sequences sampled from individuals across the state of Florida, for whom metadata, including age and county of residence, were provided and are shown in the corresponding heatmaps. Cluster status for each external branch according to MicrobeTrace (19) and dynamic prediction using the *DeepDynaForecast* are also shown. Counties were categorized according to EHE prioritization. Branches are scaled in substitutions/site.

Clustering models, including *DeepDynaForecast* described here, assume the majority of sampled individuals within an infected population transmit at a constant, or static, rate, with a minority of individuals demonstrating transmission growth or decay, relative to the majority population. Similarly, *DeepDynaForecast* was trained on outbreak scenarios wherein dynamic risk groups comprised the minority, explaining the estimated *<* 40% of sampled Florida outbreak individuals classified as predictive of future growth or decline in transmission. Among the previously identified clustered sequences from this population, approximately 92% were considered to be associated with a pattern in the tree predictive of future growth relative to the majority of sampled individuals (Figure 5A), indicating the majority of the clusters were associated with high-risk transmission groups.

**Fig 5.**
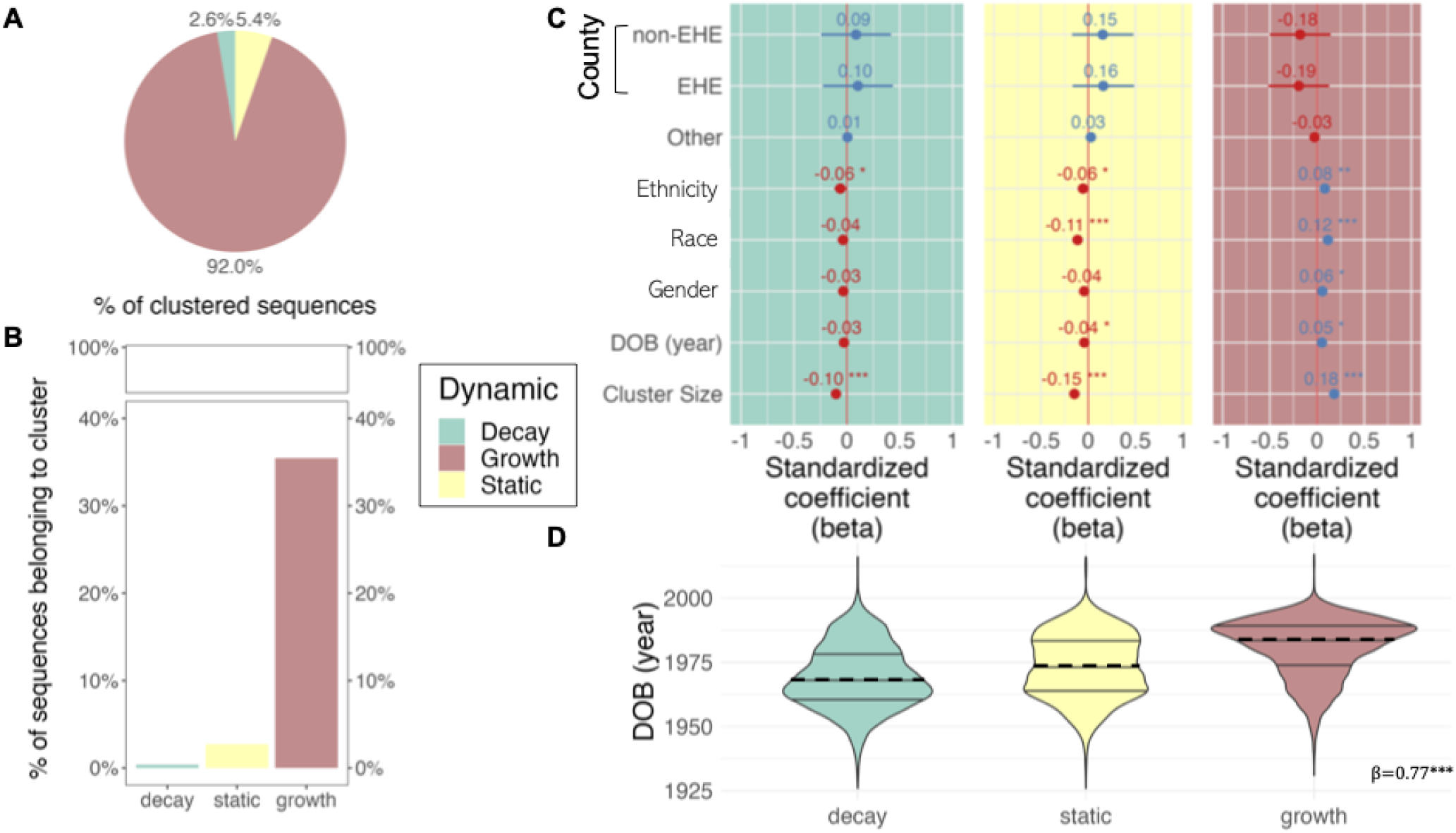
Relationship of predicted transmission dynamics with genetic clustering and patient risk factors. Transmission clusters were previously identified in Rich *et al*. (19) using MicrobeTrace (27) and the corresponding information used herein to determine the percentage of clustered sequences classified according to prediction category (A), as well as the percentage of predicted categories represented by clustered sequences (B). A multivariate logistic regression model was used to identify predictors for each transmission category, regardless of clustering status (C). Only risk factor categories with significant or marginally significant predictors are depicted, with the exception of the inclusion of county of origin, owing to its importance in clustering (28). A linear regression model, treating prediction categories as ordinal, was used to quantify the relationship between year of birth of individuals and prediction status (D). Prediction categories refer to growth, decay, or static transmission relative to the background, or majority, infected population and were determined using the trained PGLSTM model implemented in *DeepDynaForecast*.

However, *<* 35% of leaves within the tree associated with future growth were previously designated as clustered sequences (Figure 5B), indicating the majority of sequences potentially contributing to increased transmission have no clear epidemiological links. Even lower percentages (*<* 2%) were observed for lineages associated with slowing, or decaying, transmission, suggesting transmission cluster identification is not necessarily sufficient for identifying dynamic transmission patterns and high-priority groups for public health interventions. We, therefore, applied a multivariate, logistic regression model to determine if available risk factors could explain transmission dynamics, regardless of cluster affiliation. Despite prioritization based on incidence, EHE county status was highly mixed within the phylogeny (Figures 4) and not considered a significant predictor for transmission dynamics (Figure 5C). Cluster size was significantly associated (*p <* 2.5*E −*07) with transmission dynamics -individuals within previously identified larger clusters (*>* 11 individuals) tended to be predicted as contributing to transmission growth, whereas predicted static and decaying transmission were associated with smaller clusters. Race/ethnicity was considered a significant predictor (*p <* 0.05) for all three dynamic statuses, with two racial/ethnic groups exhibiting strong positive association with growth, negative association with static transmission, and one group negatively associated with decaying transmission. Gender demonstrated similar association patterns, though not considered significant for decaying transmission. Year of birth (YOB) was also considered a significant predictor (*p <* 0.05) for static and growing transmission, increasing in significance when considered in the context of all dynamic prediction categories in an ordinal regression model (*β* = 0.77, *p* = 0) (Figure 5D). Individuals over 55 years of age (median) were predicted to contribute to slowed transmission, whereas transmission growth was prominent among individuals under the age of approximately 39 years (Figure 5D).

## Discussion

*DeepDynaForecast* is a method based on graph neural network (GNN) principles capable of learning subtle topological patterns within a Phylogenetic tree structure and their association with underlying transmission dynamics. When applied to outbreaks representing both acute and chronic pathogenic infections, *DeepDynaForecast* predicted with 88.1% accuracy the dynamics of transmission clusters within *>* 15, 000 simulated outbreak phylogenies, where 90.7% and 88.9% of lineages contributing to transmission growth and decline, respectively, were identified successfully. Training of the model using stochastic transmission rates and mixed infection types enhanced not only overall performance in comprehensive scenarios but also performance when applied to individual outbreaks separately. These results indicate that *DeepDynaForecast* is a promising tool. As empirical data becomes increasingly available, it can learn and predict more diverse and comprehensive scenarios.

When applied to an HIV outbreak in the state of Florida from 2012-2017, the predictive model both corroborated previous studies but also alluded to a more informative method compared to risk group identification approaches through genetic clustering. Clustering approaches, such as HIV-TRACE (7) and MicrobeTrace (27), rely on pre-determined genetic distance thresholds and cannot discriminate groups that represent an increasing risk of transmission from those that show signs of waning transmission. Existing dynamic forecasting methods also have limitations. Some focus only on forecasting growth dynamics (29), others neglect to incorporate individual hierarchical information (18), and some rely on preidentified clusters (22). Alternatively, *DeepDynaForecast* relies solely on the data contained within the given phylogeny to identify individuals that exhibit the potential for future transmission that is elevated (e.g., increased *R*0) or slowed, or even transmission that is likely to increase in rate deterministically (i.e., continuing to increase). Moreover, identification of genetic clusters is not required. In the example of the HIV outbreak, the relatively high prevalence of classified “growing” lineages/nodes among previously identified clusters as part of the HIV outbreak is indicative of groups for whom transmission is largely uncontrolled through existing public health approaches. However, as the majority of sequences classified as “growing” according to the model had no clear epidemiological links (unclustered), the utility of transmission cluster identification in identifying high-priority groups for public health interventions may need to be re-evaluated. Even lower percentages (*<* 2%) of nodes classified as decaying, or slowing, transmission were associated with clustered individuals, indicating that interventions may be working for certain populations, but our knowledge of these groups is limited by our inability to detect them with existing clustering approaches. The associations of age, racial/ethnic group, and gender confirm prior findings and highlight the existence of disproportionate treatment allocation, potentially unrelated to geographical residency within Florida, despite prioritization of specific counties according to HIV incidence as part of the End HIV Epidemic program, as these counties collectively were not considered a significant predictor for transmission dynamics. We elected not to provide the effect ratio among sociodemographic groups because it could be subject by confounding or selection bias, which is not possible to determine within the current framework. We recognize the importance of surveillance data for informing public health policy (30, 31), and we believe that in the principle of beneficence there is a tangible public health benefit in performing research using surveillance data, but we also believe that it is of utmost importance to report results to avoid misinterpretation and risk of harm.

## Methods

### Simulation of dynamic transmission clusters

Simulation of early-midway epidemic outbreaks was performed using the *nosoi* (20) agent-based stochastic simulation platform, which is designed to take into account the influence of multiple variables on the transmission process (e.g., population structure and dynamics) to create complex epidemiological simulations. Beginning with a single infected individual, rate of transmission of infection to susceptible individuals varied according to two scenarios -1) transmission of an acute infectious respiratory virus (ARI) or 2) transmission of *Mycobacterium tuberculosis* (TB) among populations of HIV-infected or HIV-uninfected individuals, as described in (32). Specific paramaters for each outbreak scenario are described in more depth below.

### Simulation of a structured ARI outbreak

For the simulated ARI outbreak, rate of transmission of infection was dependent on the following: 1) the time since initial infection, 2) duration of infection, 3) number contacts, 4) infection rate, and 5) the risk group to which the susceptible individual belongs (Supplementary Table 1). Incubation periods were specified following infection, during which infected individuals were not permitted to transmit. Individuals were removed from the simulation after an infectious period of ∼14 days. The number of contacts per infected individual varied according to risk group and number of individuals infected within that group so as to allow for varying dynamics across risk groups. Infection rate was also allowed to vary according to risk group. The rate of infection from a background individual was 1.8 *×* 10^*−*3^ after at least two background individuals had been infected at the start of the simulation. This rate was used to ensure clusters did not exceed in number the background population. The rate of specific infection of dynamic risk groups was two-fold relative to static groups, owing to their lower representation (2:4). Following initiation, transmission was isolated to the risk group (i.e., probability of zero for infecting an individual in another cluster or background population). The number of contacts for groups A-E were picked from a normal distribution with group-specific means and standard deviation, representing clusters for which the rate of secondary infection (*R*_*e*_) remained steady, or static. The number of contacts for F and G, however, were derived from the following linear function:

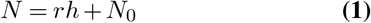

where *N*_0_ is the initial number of contacts, and *r* is the rate of change dependent on the current number of actively infected hosts in the simulation (*h*). Cluster F was considered to be experiencing an increasing rate of growth in transmission over time, whereas G was considered to be decaying over time (Supplementary Table 1). Multiple static clusters were also incorporated with varying contact parameters in order to determine indirectly the relationship of these parameters with branching patterns and thus influence on cluster classification. Each of a total of 10, 000 simulations was run for 365 days or until a total of 10, 000 hosts were infected.

### Simulation of a structured TB/HIV outbreak

A previously described co-infection outbreak model describing the impacts of HIV infection on the spread of TB was adapted herein from Goldstein *et al*. (33). Briefly, the mean incubation period, though more appropriately referred to as the latent period for TB, was 9 months among hosts that became infectious, and the mean infectious period was 3 months. The number of contacts per infected individual varied according to risk group and number of individuals infected within that group so as to allow for varying dynamics across risk groups. Infection rate was also allowed to vary according to risk group, and, unlike the ARI outbreak, transmission to individuals outside of each risk group was permitted. The rate of infection from a background individual was 1.8 ×10^*−*2^ after at least two background individuals had been infected at the start of the simulation, and the relative rate of infection of an individual belonging to a particular risk group was dependent on the risk group -two main risk groups were included, representing TB-infected individuals living with or without HIV. These groups were further split into static and dynamic transmitting groups. Based on a reported average of 53% percent of infection recipients harboring HIV (33), infection rate for HIV risk groups A and D were 0.53*/*2 = 26.5%. Remaining non-HIV-infected risk groups B and E comprised the remaining 47% (equally represented). The number of contacts for groups A and B were picked from a normal distribution with group-specific means and standard deviation, representing clusters for which the rate of secondary infection (*R*_*e*_) remained steady, or static. The number of contacts for F and G, however, were derived from the linear equation above and are also described in Supplementary Table 1. Group F was considered to be experiencing an increasing rate of growth in transmission over time, whereas G was considered to be decaying over time (Supplementary Table 1). Group B was thus considered to exhibit static transmission. Each of a total of 10, 000 simulations was run for 8 years or until a total of 10, 000 hosts were infected.

### Reconstruction of sampled transmission tree from simulated outbreaks

Recipients were only sampled during their infectious periods, with the sampling time equally likely at any point in this time frame. Only simulations where at least 50 individuals were infected were accepted. One representative clade within the background population was chosen at random from the corresponding internal nodes and maintained all individuals (5-100), representing true clusters of direct transmission, or transmission chains. Remaining risk groups were sampled randomly, ranging in frequency from 20-100% of the original cluster population. The background population was also downsampled at a frequency of 20%, representing a more realistic surveillance scenario. Hosts not included within this sample were pruned from the full tree to obtain the final set of simulated trees used for tree statistic calculation and deep learning models. A relaxed molecular clock (evolutionary rate in time) was assumed, and branch lengths were scaled in time (substitutions/site/year) using a uniform distribution rate multiplier (∼ *𝒰* (8 ×10^*−*4^, 0.001)), allowing for both genetic distance and time to be used as distinct weights in the neural network models.

In this study, 10, 000 simulations beginning with a single infected individual were performed on ARI and TB individually, resulting in 8, 574 and 7, 182 successful outbreaks, respectively. In ARI simulations, Phylogenetic trees include 33, 713, 775 nodes. Nodes are in three main categories -static, growing, or decaying -based on pre-defined transmission dynamics in the simulation (Supplementary Table 1), resulting in 31, 920, 116 static, 589, 119 decay and 1, 204, 540 growth samples. For TB simulations, 7, 182 trees include 21, 417, 772 edges and 21, 424, 954 nodes with 20, 884, 036 static, 51, 419 decay, and 489, 499 growth samples. Tree files can be downloaded from the Github repository (https://github.com/lab-smile/DeepDynaForcast).

### Simulation data pre-processing and augmentation

Due to the highly unbalanced input distribution among the three classification categories (see Supplementary Figures 3 and 5 for details), the data represents a realistic scenario in which sampling is biased according to prevalence within the population. Nevertheless, the data was redistributed for model development using a tree-based random split method with a 60%-20%-20% ratio. This approach ensures that all nodes of a single Phylogenetic tree are only allowed to be split into either the training, validation, or test subset with the proposed probabilities, preventing data leakage. Before feeding the data into the learning algorithms, we performed data pre-processing for raw inputs. For edge features (See Supplementary Tables 2 and 3), we first applied an inverse hyperbolic sine (ArcSinh) transformation, approximating the natural logarithm of raw values while retaining zero-valued observations, followed by z-score normalization. Due to the difference on the unit of ARI and TB trees, the normalization was performed on each dataset individually. Node features were not available from the simulations but are necessary for *DeepDynaTree* modeling. Therefore, we randomly initialized features of size 16 during the training process to improve model robustness and used fixed zero vectors when testing.

In accordance with our previous research (22), graph neural network models are limited to emerged nodes in transmission clusters. To enhance the model’s performance in the early stages of outbreaks with small clusters, we utilized a data augmentation strategy for the training data. In each generated cluster, branches were chopped at a random chopping rate ranging from 0 −100%, prioritizing nodes close to the simulation’s end. This downsizes the number of nodes in each cluster, allowing the model to focus on forecasting small clusters. The distributions of edge features after preprocessing and data augmentation are summarized in Supplementary Tables 2 and 3 and Supplementary Figures 2 and 4.

### DeepDynaForecast Modeling

*DeepDynaForecast* is developed to utilize the topological information and branching patterns within the phylogeny, including genetic distances, to predict the dynamics of external nodes successfully. The Phylogenetic tree is modeled as a bi-directed graph, and our model predicts the dynamic status of leaves. The following sections introduce two major components -primal-dual learning structure and cross-layer predictions.

### Primal-dual graph LSTM module (PDGLSTM)

The PDGLSTM (21, 22) constructs a dual graph (i.e., line graph) of the Phylogenetic tree, where the nodes and edges correspond to the tree’s branches and nodes respectively. The PDGLSTM processes both the original (primal) and dual graphs, learning node and edge representations concurrently by applying Graph LSTM (GLSTM) models.

The GLSTM integrates a Long Short-Term Memory (LSTM) model to learn sequential representations that emerge from several message-passing iterations. As depicted in Figure 1, each edge and node aggregates the representations of adjacent nodes or edges. The neural networks *ϕ*_*E*_ and *ϕ*_*N*_ encode these aggregates into low-dimensional messages. These messages are then fed into their corresponding LSTM modules to facilitate the update of node and edge representations.

Formally, given the node representation 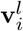 of node *i* at *l*-th layer and its aggregated message **m**_*i*_, the node representation is updated according to the LSTM updating rule:

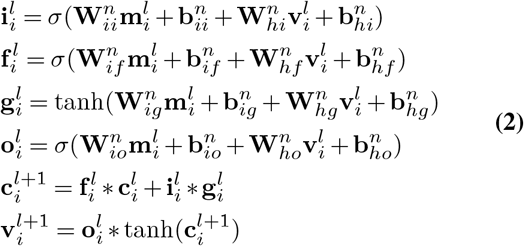

In these equations, 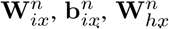, and 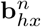 are model parameters for *x ∈ {i, f, g, o}*. 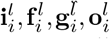, represent the outputs of input gate, forget gate, memory cell and output gate, and *σ*(*·*) denotes sigmoid function. Both **c**_*i*_ and **v**_*i*_ are randomly initialized. Similarly, the edge representation 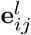 at layer *l* is updated via an Edge-LSTM model:

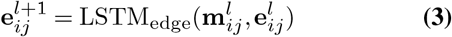

where the **m**_*ij*_ represents the aggregated message for edge *e*_*ij*_ and the detailed computation of *LST M* is as same as Eq. 2, except replacing the node inputs with the edge and its associated message information. In our setting, both Node-LSTM and Edge-LSTM models are configured to have hidden sizes of 128.

Low-dimensional messages are derived by aggregating the representations of adjacent nodes and edges. For example, for node *i*, the aggregated message **m**_*i*_ is created by applying the neural network *ϕ*_*N*_ to the representations of inbound edges and the node itself:

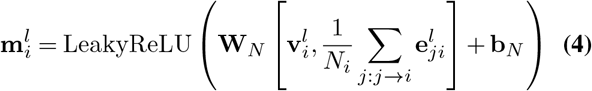

Here, *j →i* symbolizes all inbound edges to node *v*_*i*_, and [·, *·*] signifies vector concatenation. **W** and **b** are learnable parameters, mapping the concatenated vector from 2 ∗128 = 256 dimensions to a 16-dimensional space.

Similarly, the message information **m**_*ij*_ for edge *e*_*ij*_ is generated by *ϕ*_*E*_:

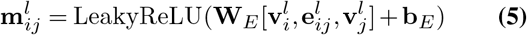

This is achieved by using the representations of edge *e*_*ij*_ and its neighboring nodes *i* and *j*.

### Cross-layer Prediction module (CLP) and dynamic prediction

Upon completing *L* message-passing iterations, the Cross-layer Prediction (CLP) module predicts the dynamics of leaf nodes while masking the internal nodes during this prediction phase. A series of neural networks, denoted as *ψ*_*L*_, are utilized to predict the leaf node dynamics. These networks use various levels of node representations, **v**. To enhance the robustness of the final result, this process is accompanied by dropout layers and summation operations:

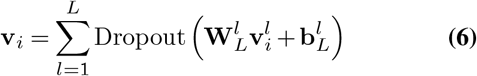

In this equation, parameters 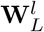 and 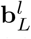 are associated with the network 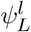 for the node representation at the *l*-th layer. *L* represents the number of message-passing iterations.

The PDGLSTM accumulates extensive information from multi-hop neighboring nodes and edges through iterative application of the primal-dual message passing and state updating steps. This process results in an enhanced, highly informative node representation for the final prediction. In our configuration, the message-passing steps were repeated 20 times. This means the receptive field of each node spans up to 20-hop neighboring nodes and edges. The consolidated node representation, **v**_*i*_, is subsequently processed by a softmax layer to produce the final prediction.

### Model evaluation metrics

In this study, our proposed DeepDynaForecast was comprehensively evaluated on six performance metrics, including balanced accuracy, precision, F1-score, area under the receiver operator characteristic curve (AUROC), Brier score (BS), and Cross-entropy (CE). Precision, F1-score, and AUROC are broadly used measures of the discriminability of a pair of classes. To deal with the multi-class case, we first generated the metric values for each class in a one-vs-rest manner and then averaged them with equal weights to give the same importance to each class. Considering the highly unbalanced label distribution in our data set, equally averaging the metrics over all classes instead of weighted averaging by the reference class’s prevalence in the data can effectively avoid overestimating the models that only perform well on the common classes while performing pooling on the rare classes. This equally averaging strategy is also named macro-average. The balanced accuracy is defined as the average of recall obtained on each class (34). Besides the above metrics calculated based on the “hard” classification results (i.e., final category information only), we also measured the BS and CE to evaluate the models’ “soft” predictions, wherein the metrics were calculated on the prediction probabilities. Eq. 7 illustrates the definition of weighted BS and weighted CE values,

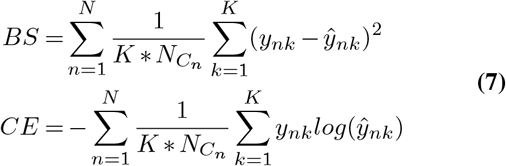

where *ŷ*_*nk*_ denotes the predicted probability of sample *n* for label *k*, and *y*_*nk*_ is the ground truth with binary value. *C*_*n*_ represents the class of sample *n. N* is the number of samples in the data set and *K* is the number of classes where it equals to 3 in our case. Lower BS and CE values thus indicate better prediction performance.

### GCN and GIN baseline models

We selected Graph Convolutional Network (GCN) (23) and Graph Isomorphism Network (GIN) (15) as two baseline models in the dynamic prediction task and trained them on combination of ARI and TB datasets. To ensure a fair comparison, we configured the number of layers to be 20 and set the hidden size at 128. These configurations align with the hyperparameters used in the *DeepDynaForecast*. For GIN, we employed summation as the aggregation type and initiated the *ϵ* value at 0.

### Visualizing node representations with UMAP Projection

We used the Uniform Manifold Approximation and Projection (UMAP) (24) 2D projection to visualize the node representations learned from the *DeepDynaForecast* architecture. We concatenated the hidden representation of nodes from each message-passing iteration into a vector with a size of 20 ∗128 = 2560 and selected the top-100 principal components to feed into the UMAP calculation. For the GCN and GIN baseline models, we employed the node representations from just before the last classification layer in the UMAP projection calculations.

### Implementation Details

This section introduces the implementation details of *DeepDynaForecast* models. A weighted Cross-entropy loss with inversely proportional to class frequency weights was applied for model optimization. Hyper-parameters of all the methods were extensively optimized on the same validation set for a fair comparison. The model took approximately 1 day to train for around 100 epochs with mini-batch of size 4. An adam optimizer was used for training with an initial learning rate 1 ×10^*−*3^, and 90% reduction was applied to the learning rate if the validation loss did not improve in the consecutive 50 epoches until reaching the the minimum value 1 ×10^*−*4^. We finetuned all the hyper-parameters on the validation data set. All experiments were performed on a workstation with 12 Intel Core i7-5930K CPUs and a single Nvidia GeForce GTX TITAN X GPU card.

## Code availability

The DeepDynaForecast model code, along with all other code written to generate the results shown in this study, is available on Github under MIT Licence at: https://github.com/lab-smile/DeepDynaForecast.

## Data availability and ethics statement

The authors abide to the Declaration of Helsinki. The study protocol was approved by the University of Florida’s Institutional Review Board (IRB) and by FDOH’s IRB (protocol no. IRB201901041 and 2020-069, respectively) as exempt. We received HIV sequence data and metadata extracts from FDOH in a fully de-identified format compliant to the Health Insurance Portability and Accountability Act (HIPAA). For replication purposes, data request to the FDOH can be made according to state, federal regulations and compliance with required ethical and privacy policies, including IRB approval by FDOH and execution of data user agreements. Requests are independently reviewed by FDOH.

## ACKNOWLEDGEMENTS

This project was supported by the US National Institute of Allergy and Infectious Diseases (R01AI145552), and the Stephany W. Holloway University Chair in AIDS Research. The funders had no role in the writing of the manuscript or the decision to submit it for publication. The findings and conclusions in this report are those of the authors and do not necessarily represent the views of the FDOH or the other funders.

## Supplementary Material

**Table 1.** Simulation information for each outbreak and risk group.

| Outbreak scenario | Risk group | Contact parameters<sup>1</sup> | Mean P(T)<sup>2</sup> | Mean $R_0$ |
| --- | --- | --- | --- | --- |
| ARI | A | $\sim \mathcal{N}(16, 1)$ | 0.015 | 2.2 |
| ARI | B | $\sim \mathcal{N}(4, 1)$ | 0.09 | 3.2 |
| ARI | C | $\sim \mathcal{N}(4, 1)$ | 0.11 | 3.9 |
| ARI | D | $\sim \mathcal{N}(6, 1)$ | 0.09 | 4.9 |
| ARI | E | $\sim \mathcal{N}(4, 1)$ | 0.12 | 4.3 |
| ARI | F | $N_0 = \sim \mathcal{N}(4, 1),$<br>$r = \sim \mathcal{N}(0.018, 0.009)$ | 0.11 | 0.8 |
| ARI | G | $N_0 = \sim \mathcal{N}(6, 1),$<br>$r = \sim \mathcal{N}(0.18, 0.09)$ | 0.11 | 9.8 |
| TB/HIV | A | $\sim \mathcal{N}(20, 5)$ | 0.025 | 1.5 |
| TB/HIV | B | $\sim \mathcal{N}(20, 5)$ | 0.015 | 0.9 |
| TB/HIV | F | $N_0 = \sim \mathcal{N}(15, 1),$<br>$r = \sim \mathcal{N}(0.18, 0.09)$ | 0.015 | 0.7 |
| TB/HIV | G | $N_0 = \sim \mathcal{N}(20, 1),$<br>$r = \sim \mathcal{N}(1.8, 0.9)$ | 0.025 | 1.5 |

<sup>1</sup> Single values represent mean number of contacts for static clusters.<sup>2</sup> Probability of transmission**Table 2. Summary statistics for edge features in ARI.**

**Table 2.** Summary statistics for edge features in ARI.

| Features | Min | Max | Mean | Std |
| --- | --- | --- | --- | --- |
| Time (days) | 0.000 | $8.600 \times 10^1$ | 7.771 | 6.221 |
| Genetic distance (substitutions/site) | 0.000 | $8.072 \times 10^{-2}$ | $6.994 \times 10^{-3}$ | $5.629 \times 10^{-3}$ |

**Table 3.** Summary statistics for edge features in TB.

| Features | Min | Max | Mean | Std |
| --- | --- | --- | --- | --- |
| Time (weeks) | 0.000 | $1.940 \times 10^2$ | 9.470 | 8.067 |
| Genetic distance (substitutions/site) | 0.000 | $1.834 \times 10^{-1}$ | $8.524 \times 10^{-3}$ | $7.296 \times 10^{-3}$ |

**Table 4.** Performance for DDF_ARI_TB. We evaluated the performance of DDF_ARI_TB on different leaf groups, including background leaves only (Background leaves), leaves identified as high-risk clusters (Cluster leaves), and a combination of them (All leaves). For cluster leaves and all leaves, to mitigate the impact of unbalanced label distribution, metrics—including accuracy, F1-score, precision, and area under the receiver operating characteristic (AUROC)—were uniformly aggregated across the three classes. Weighted Brier score (BS) and weighted Cross-entropy (CE) were calculated based on predicted probabilities adjusted by the inverse prevalence of classes, providing “soft” evaluations of the models. As all background leaves are static, only accuracy, BS, and CE are shown here. For each testing dataset, models were assessed on ARI trees, TB trees, and a combination of both.

| Dataset | Leaf group | Balanced Accuracy ↑ | Macro F1 ↑ | Macro Precision ↑ | Macro AUROC ↑ | Weighted BS ↓ | Weighted CE ↓ |
| --- | --- | --- | --- | --- | --- | --- | --- |
| ARI | Background leaves | 0.875 | — | — | — | 0.184 | 0.320 |
|  | Cluster leaves | 0.724 | 0.549 | 0.606 | 0.869 | 0.442 | 0.797 |
|  | All leaves | 0.897 | 0.494 | 0.451 | 0.969 | 0.155 | 0.279 |
| TB | Background leaves | 0.883 | — | — | — | 0.180 | 0.336 |
|  | Cluster leaves | 0.590 | 0.417 | 0.460 | 0.806 | 0.551 | 0.935 |
|  | All leaves | 0.744 | 0.392 | 0.381 | 0.925 | 0.345 | 0.604 |
| ARI+TB | Background leaves | 0.878 | — | — | — | 0.182 | 0.327 |
|  | Cluster leaves | 0.711 | 0.530 | 0.572 | 0.859 | 0.443 | 0.786 |
|  | All leaves | 0.881 | 0.459 | 0.424 | 0.964 | 0.175 | 0.315 |

**Fig. 1.**
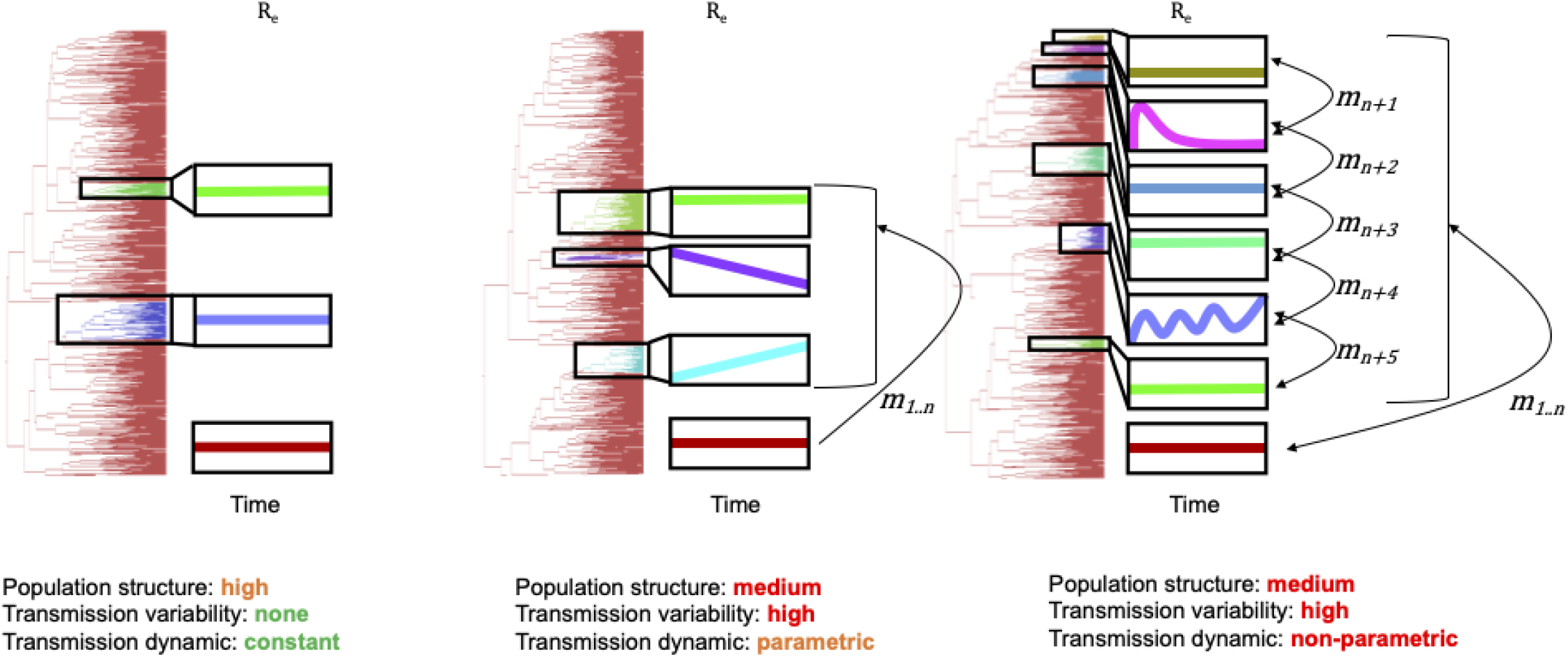
Ranging complexity of tree topological features resulting from a structured infected population. *n* transmission clusters, distinct from the background population (maroon), can vary in effective reproductive number (*R*_*e*_) over time (x-axis) and rate of infection (*m*) by or to individuals from other groups. Variations in transmission dynamics are imprinted in branch patterns within the corresponding phylogeny and can aid in identifying groups of interest.

**Fig. 2.**
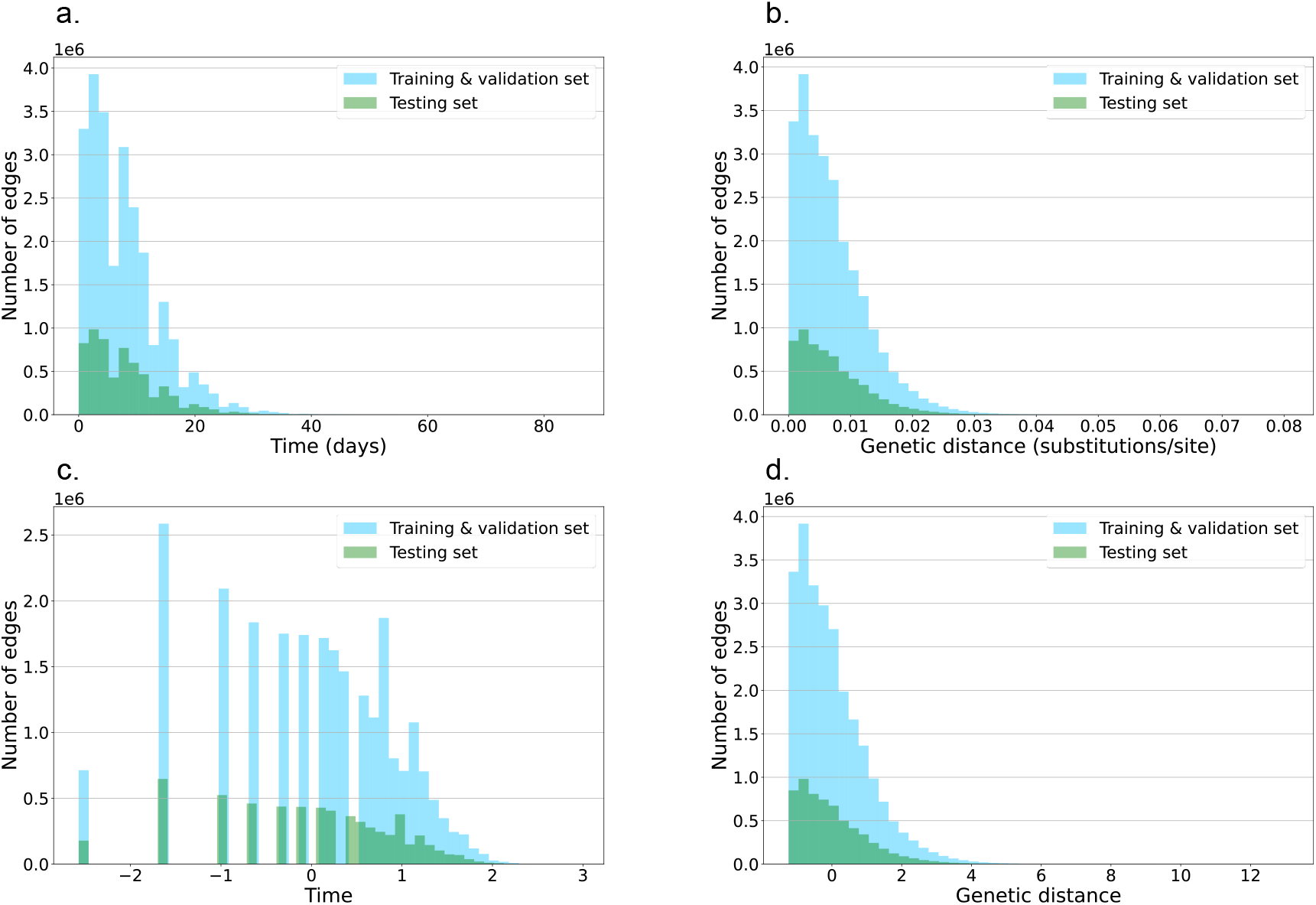
Distribution of the edge features on ARI simulations. **a**, and **b**, are distributions of the raw edge features: time and genetic distance. **c**, and **d**, are distributions of the edge features processed by an ArcSinh transformation and a z-score normalization.

**Fig. 3.**
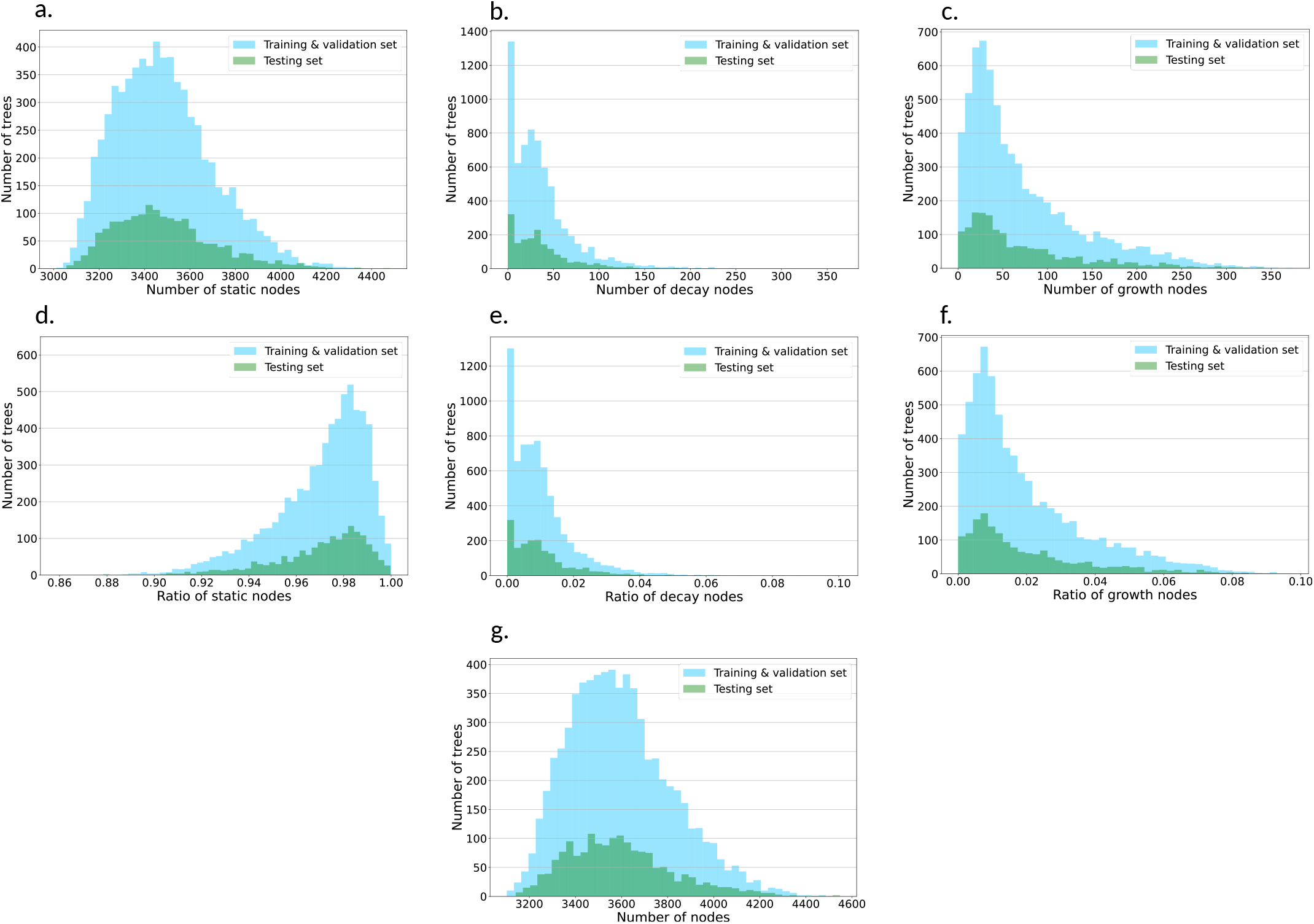
Label distribution on ARI simulations. **a-c** are histogram of the number of static, decay, and growth nodes among the trees, and **d-f** show the distribution of classes’ ratio. Plot **g** is the histogram of the number of nodes on the trees.

**Fig. 4.**
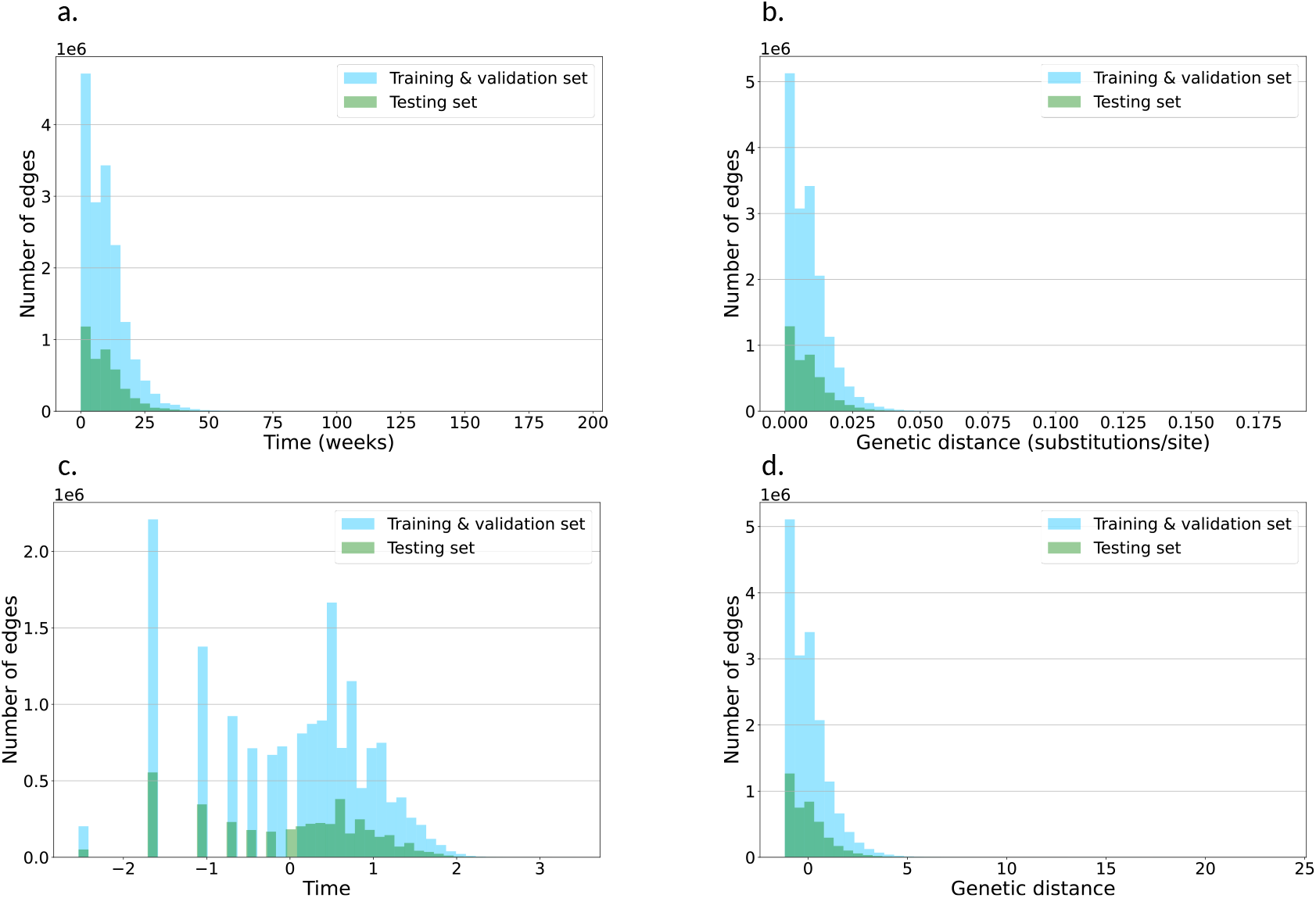
Distribution of the edge features on TB simulations. **a**, and **b**, are distributions of the raw edge features: time and genetic distance. **c**, and **d**, are distributions of the edge features processed by an ArcSinh transformation and a z-score normalization.

**Fig. 5.**
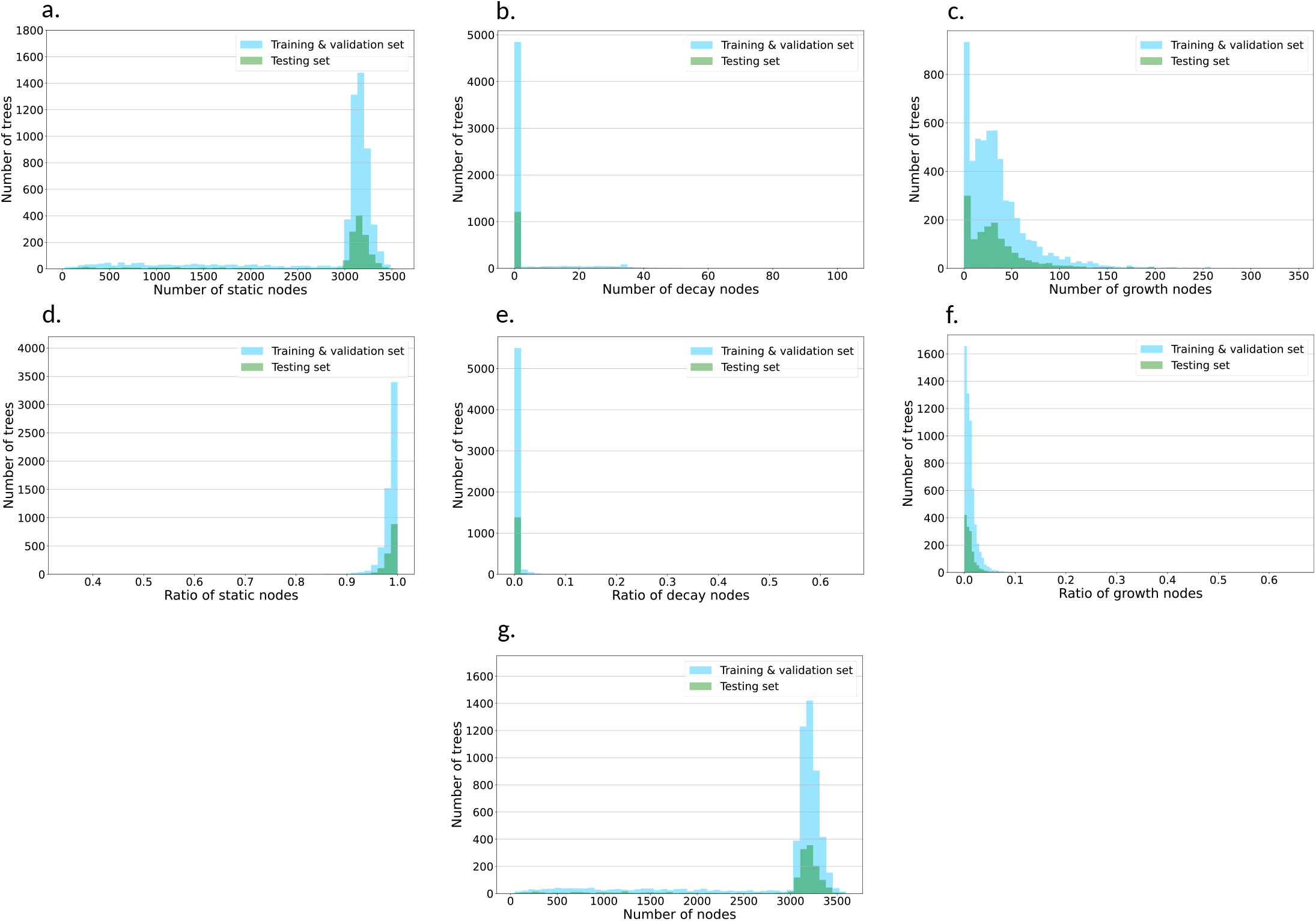
Label distribution on TB simulations. **a-c** are histogram of the number of static, decay, and growth nodes among the trees, and **d-f** show the distribution of classes’ ratio. Plot **g** is the histogram of the number of nodes on the trees.

## Notes

### Competing Interest Statement

The authors have declared no competing interest.

